# RNA2seg: a generalist model for cell segmentation in image-based spatial transcriptomics

**DOI:** 10.1101/2025.03.03.641259

**Authors:** Thomas Defard, Alice Blondel, Sebastien Bellow, Anthony Coleon, Guilherme Dias de Melo, Thomas Walter, Florian Mueller

**Affiliations:** Center for Computational Biology, Mines Paris PSL, Boulevard Saint Michel, Paris, 75017, France.; Institut Curie, PSL University, 26 rue d’Ulm, Paris Cedex, 75248, France.; U900, INSERM, 26 rue d’Ulm, Paris Cedex, 75248, France.; Institut Pasteur, Université Paris Cité, Centre de Ressources et Recherches Technologiques (UTechS-PBI, C2RT, Paris, France.; Institut Pasteur, Université Paris Cité, scBiomarkers UTechS, Paris, France.; Institut Pasteur, Université Paris Cité, Lyssavirus Epidemiology and Neuropathology Unit, Paris, France.

**Keywords:** Spatial Transcriptomics, Cell Segmentation, Deep Learning

## Abstract

Imaging-based spatial transcriptomics (IST) enables high-resolution spatial mapping of RNA species. A key challenge in IST is accurate cell segmentation to assign each RNA molecule to the right cell. Here, we present RNA2seg, a novel segmentation algorithm trained on over 4 million cells from MERFISH and CosMx datasets across seven organs using a teacher-student training scheme. RNA2seg integrates RNA point clouds and all available membrane and nuclear stainings. Validation on manually annotated data shows superior performance including in zero-shot and few-shot settings. The method is available as a documented pip package: https://github.com/fish-quant/rna2seg.

## 1 Background

Understanding the spatial organization of tissues at the single-cell level is essential for advancing our knowledge of disease progression and development [1, 2]. Advances in spatial transcriptomics (ST) now allow the measurement of gene expression at cellular and even subcellular resolution, offering valuable insights into cell type distribution, cell-to-cell communication [3], and disease-specific signatures [4].

ST can be broadly categorized into sequencing-based and imaging-based spatial transcriptomics (IST). While sequencing-based techniques provide coverage of the full transcriptome [5], they do not offer cellular or subcellular resolution. In contrast, IST enables the localization of RNA molecules in their cellular environment, albeit at the cost of being limited to a predefined set of target genes [6, 7].

A critical aspect of IST for assessing single-cell gene expression (GE) and subsequent cell type identification is the accurate assignment of measured RNA to the correct cell. To achieve this, IST typically provides several staining channels, including nuclear and membrane markers, as well as non-specifically labeled transcripts. Segmentation algorithms then assign each transcript to a specific cell, generating a spatially resolved count matrix for subsequent analysis. This segmentation step is critical, as errors in RNA-cell association result in erroneous transcriptional profiles and thus affect all downstream analyses, such as cell type identification [8].

Segmentation algorithms for IST can be divided into staining-based and point cloud-based methods. The former rely on dedicated membrane or nuclei stainings and allow the application of deep learning methods such as Mesmer [9] or Cellpose [10], which can achieve performance close to human-level. However, one limitation comes from the highly variable quality of the cell staining in IST. Indeed, the fluorescent reporters in these channels often bind to cell-type specific proteins leaving the rest of the tissue unlabeled [11]. Consequently, segmentation accuracy — and therefore the accuracy of all downstream analyses — tends to vary significantly across the tissue, introducing biases and unacceptable inconsistencies in the analysis results.

On the other hand, the measured RNA signal is itself also informative on cellular boundaries. The density of RNAs varies across the cell, and the transcriptional profiles of cells of different type are per definition different. These observations can be used to design cell segmentation algorithms that operate directly on the RNA point clouds. Examples include Baysor [12], SCS [13], PciSeq [14] and ComSeg [8].

While these methods are promising alternatives in the absence of dedicated cell and nuclear staining, they come with several limitations. For instance, they often struggle to accurately separate neighboring cells with highly similar RNA compositions. Furthermore, these approaches typically start from segmented nuclear regions [8, 14] and subsequently associate RNAs with segmented nuclei. By design, they are unable to handle missing nuclei, a common issue when working with 2D tissue sections derived from a three-dimensional volume. Deep Learning models trained on RNA point clouds and DAPI images have also been proposed recently [15, 16]. However, these methods are limited by the scarcity of annotated data [15] or rely on biological prior information that are hard to obtain in practice [16]. To the best of our knowledge, no existing method fully leverages all available stainings, including membrane stainings and RNA point clouds.

Here, we propose RNA2seg, a deep-learning segmentation model designed for IST data, which combines the advantages of staining and point-cloud segmentation methods. RNA2seg takes an arbitrary number of staining channels and the RNA positions as input and trains a multiple-instance segmentation model for cell segmentation. To the best of our knowledge, this is the first method integrating all available staining channels and RNA point clouds for segmentation.

Deep Learning algorithms for segmentation typically require large annotated datasets which can be very challenging to produce and are missing for IST data. To address this bottleneck, we devised a teacher-student strategy that enables RNA2seg to be trained without the need for manual annotations by leveraging the teacher model in areas with high-quality staining. This approach allowed us to train RNA2seg on a dataset of over 4 million cells from seven different human organs and two distinct IST protocols - MERFISH [6] and CosMx [11] (see Supplementary Table 1).

We validated RNA2seg against a set of 724 manually annotated cells across 5 datasets of different organs (see Supplementary Table 2). These annotated images were witheld during training. We demonstrate that RNA2seg outperforms current state-of-the-art (SOTA) methods on these manually annotated dataset. This large and unprecedented set of manually annotated cells was created using a novel visualization approach for RNA point clouds, which color-codes RNA point clouds based on co-expression, assigning similar colors to co-expressed RNA species. This approach eases the cell annotation task in regions where membrane staining is of poor quality. We make the annotated data accessible to the scientific community at https://zenodo.org/records/14912364. As our training procedure does not rely on manual annotation, we lastly demonstrate how our model can be easily and automatically fine-tuned for use with previously unseen datasets.

RNA2seg is designed for large datasets and supports input in OME-Zarr format [17] through SpatialData [18] and is implemented on top of the SOPA framework [19]. RNA2seg is available as a fully documented Python pip package (https://github.com/fish-quant/rna2seg).

## 2 Result

### 2.1 Method overview

RNA2seg is a deep learning (DL)-based method for the segmentation of cells from Spatial Transcriptomics (ST) data that integrates an arbitrary number of channels with nuclear and membrane markers and RNA positional data (Figure 1A).

**Fig. 1.**
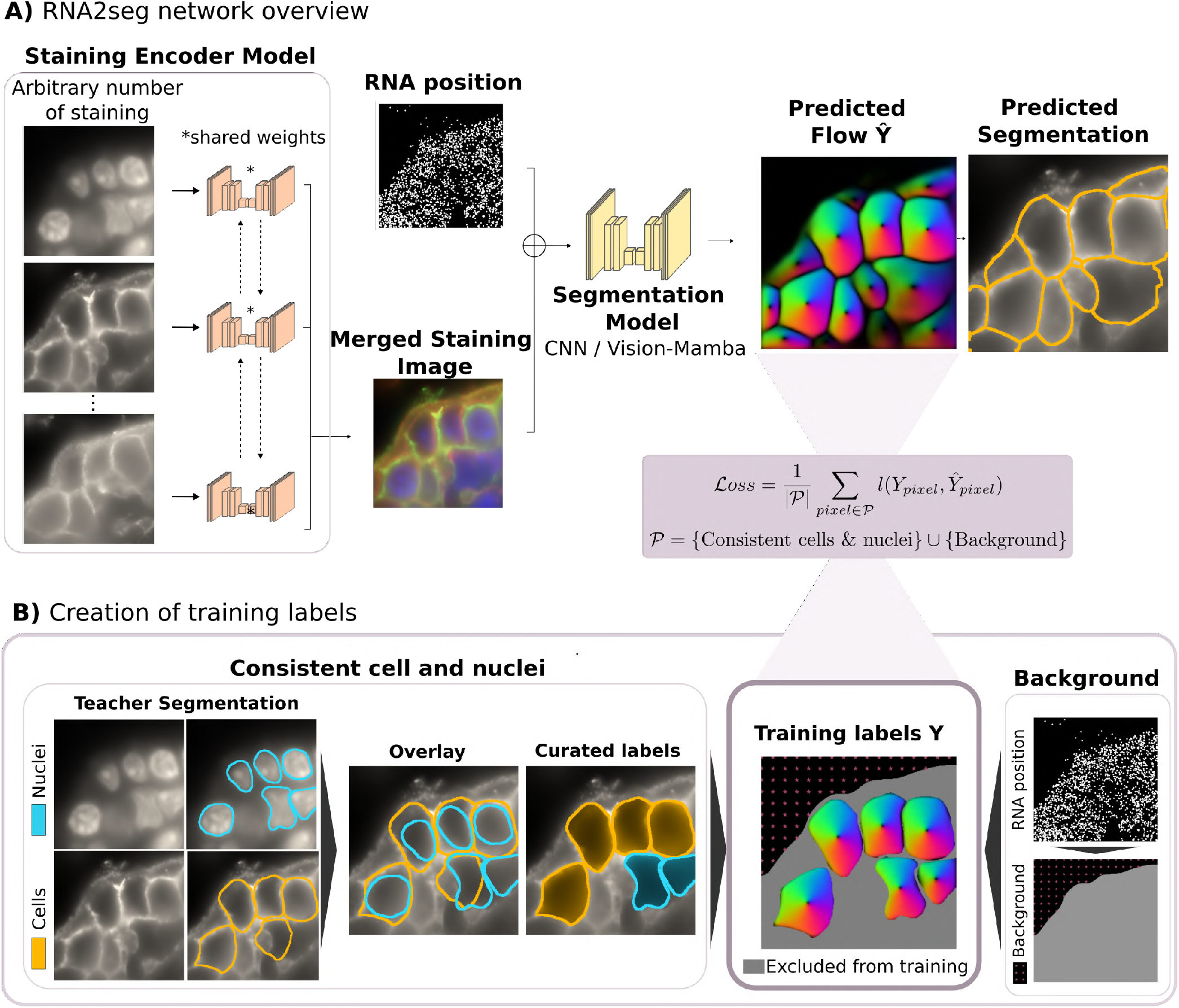
RNA2seg Methodology Overview. **A)** Segmentation model combining RNA images and multiple stainings with partial supervision. An arbitrary number of input stainings are first processed by an encoding network to produce a merged image with a fixed number of 3 channels. This encoded staining representation is then merged with an image displaying RNA positions and processed by a U-Net-like model. The final output is the flow field of the predicted cells. The model is trained on cell segmentation masks *Y* provided by the teacher network after application of consistency rules, as illustrated in B. **B)** Creation of training labels from the segmentation masks provided by the teacher network, after application of consistency rules between cell and nucleus segmentation.

To achieve this, RNA2seg merges staining images and RNA point clouds, converting RNA positions into an image. To ensure that our method is independent of the gene panels used, we assigned to each pixel the number of RNAs detected irrespective of the encoding gene, and further processed the resulting image (see Methods 5.3) to account for its sparsity. While this representation discards RNA identity and retains only positional information, we argue that the resulting network may be more easily transferable to new datasets with different gene panels unseen during training. Importantly, we also tested alternative representations comprising RNA identity without observing notable performance improvements (section 2.4).

To handle the varying number and types of nuclear and membrane staining channels, we use ChannelNet [20] - a DL framework that takes an arbitrary number of channels, extracts relevant information from them, and encodes it into a three-channel image. The merged membrane and nuclear staining representations are then concatenated with the RNA positional images to form the input of our segmentation network, which uses a U-net architecture. By incorporating information about the detected RNAs, we hypothesized that our model can achieve robust segmentation, even in regions with low-quality or missing staining.

The choice of the most efficient architecture for cell and nucleus segmentation — whether CNNs or newer, more flexible models with more parameters, such as Transformers or the recently proposed Mamba architecture — remains an active debate in the research community [21, 22]. In this study, we evaluated two architectures: CNN-based U-Net, similar to Cellpose [10], and Mamba-U-Net, a recently proposed segmentation model [23]. We refer to these models as RNA2seg-CNN and RNA2seg-Mamba, respectively. Following the Cellpose strategy, our networks are trained to predict gradient flow toward cell centers, serving as a pretext task for cell instance segmentation.

Training deep learning segmentation models typically requires large annotated datasets to achieve high performance. However, to the best of our knowledge, no large dataset with manually annotated cellular boundaries exists for image-based spatial transcriptomics data.

Manual annotation for segmentation is often time-consuming. Besides, for IST data, it is even more challenging due to several factors: the varying quality of membrane staining, the difficulty of visually interpreting RNA point-cloud data, and the scattering of relevant information across a relatively large number of channels. Consequently, the time and effort needed to generate a sufficient number of annotations for training seem prohibitive.

To address this, we developed a student-teacher framework that enables RNA2seg to be trained without manual annotations. To generate the training data for RNA2seg, we first apply a teacher pre-trained network, such as Cellpose, for cell and nucleus segmentation. Erroneously segmented cells are then filtered out, while the accurately segmented cells from the teacher model are retained, ensuring that the training process relies exclusively on reliable labels (Figure 1B). By learning the RNA spatial distribution of cells from the teacher generated labels, the model can segment even low-quality staining regions thanks to the complementary spatial RNA patterns.

To remove segmentation errors from the generated teacher labels, we rely on a set of rules (see Methods 5.2) designed to ensure consistency between nuclear and cellular segmentations. For instance, cells with multiple nuclei are removed. In regions with low staining quality, cells are often missed entirely by the teacher network. In this case, detected nuclei are used as substitutes. Of note, our training scheme for partially annotated data enables the inclusion of nuclei in the training dataset, using them as partial labels to represent cells. The underlying hypothesis being that cell and nuclear centers are relatively close, resulting in nearly equivalent prediction tasks regarding the gradient flow.

Additionally, background labels are assigned to pixels in low RNA density regions (see Methods 5.2.2). Pixels that remain unlabeled by the above procedures (i.e. neither cell nor background) are excluded from the training process through partial backpropagation [24]. The loss can thus be written as follows:

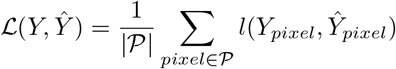

where *Y* represents the teacher segmentation, *Ŷ* is the prediction of RNA2seg, and *P* = {Curated cells & nuclei} ∪ {Background}.

Furthermore, we incorporate dropout on the membrane staining images, i.e. membrane staining image channels were set to 0 with a dropout probability of 0.25, and thus emulating missing membrane staining signals. This forces RNA2seg to learn how to segment cells solely from RNA point clouds in low quality staining areas.

Our automated student-teacher training pipeline enables large-scale model training. RNA2seg was trained on over 4 million cells across seven different human organs: breast, colon, lung, melanoma, uterine, prostate and ovarian. Two different IST technologies were included in the training set, MERFISH [6] data from Vizgen’s public Human FFPE Immuno-oncology Data Release (VHFI) and a public CosMX human lung dataset [11] (see Supplementary Table 1 for additional details). The training inputs incorporated the DAPI stain, the Poly-T stain, all of the three additional membrane staining channels available in each of these datasets, as well as the RNA positional images. The cell instance segmentations used as teacher in this study are available at: https://zenodo.org/records/14916899

### 2.2 A manually curated ground truth dataset for segmentation of IST data

Validation of cell segmentation from IST data is particularly challenging. The gold standard for validation of automatic segmentation usually involves manual annotation by experts. However, in IST, manual annotation is complicated by several factors: the quality of the membrane staining is highly variable (e.g. Figure 2B), and manual annotation of the membrane channels can therefore not be achieved in all parts of the image. Furthermore, it is difficult to visually interpret RNA point cloud data and to manually infer the correct cellular boundaries. Third, the relevant information is scattered across a relatively large number of channels. All of these factors make manual annotation of IST data particularly difficult.

**Fig. 2.**
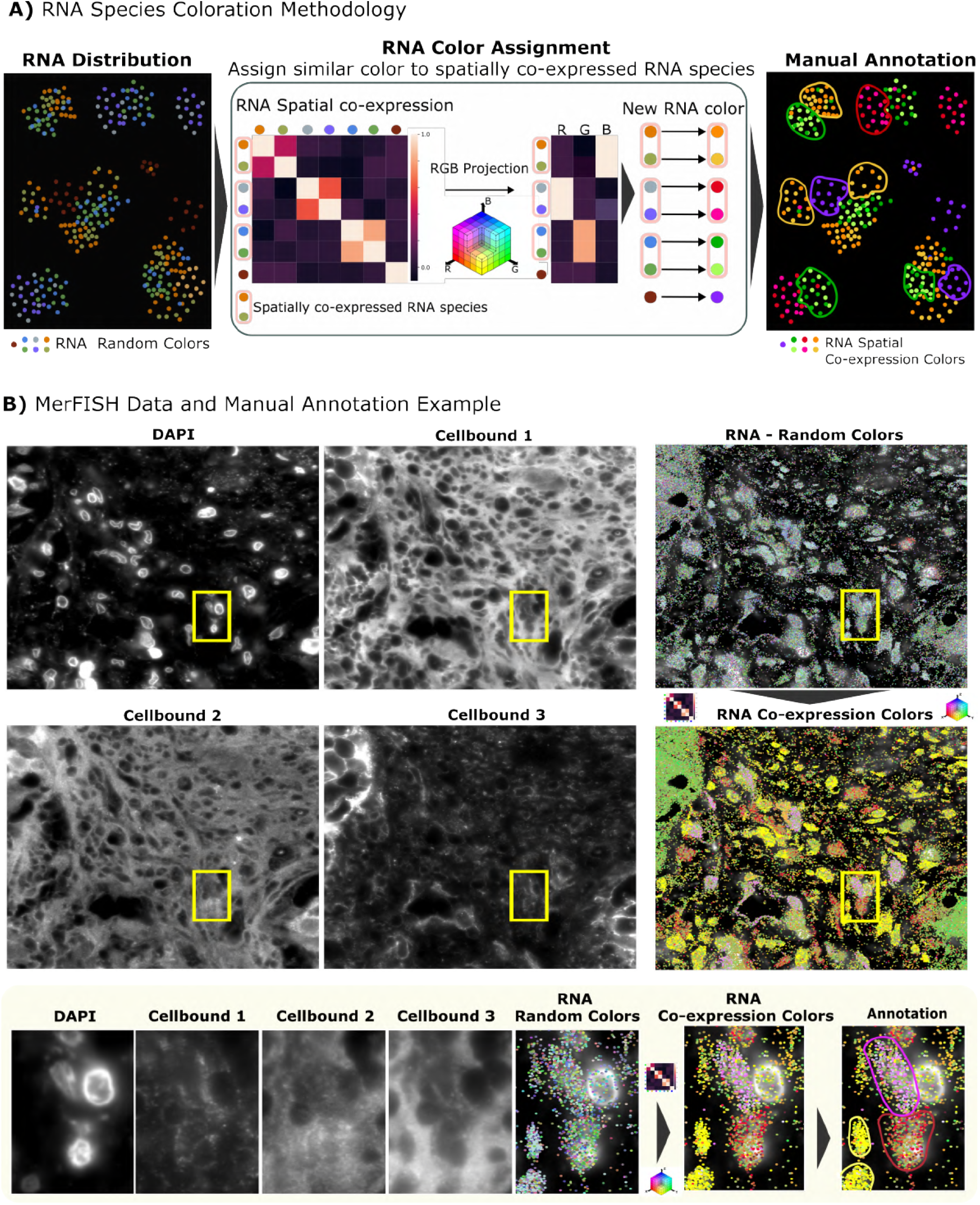
Methodology for RNA species colorization to ease manual cell annotation. **A)** Visualization with co-expression for manual annotation: The first panel shows a synthetic example of RNA distribution, where random colors are initially assigned to each RNA species. A spatial correlation matrix is then computed to quantify the recurrent neighbors of each RNA species. This matrix is projected into an RGB color space, assigning similar colors to RNA species that frequently occur as neighbors. This new color assignment simplifies manual cell annotation. **B)** Example of MERFISH staining: The top panels show MERFISH staining on VHFI breast tissue, where staining quality is heterogeneous. The bottom panels display a zoomed-in region where the staining quality is insufficient for manual annotation. However, the RNA distribution, combined with our correlation-based color visualization, enables to identify more cell contours.

For this reason, various validation strategies and quantitative metrics have been proposed that do not rely on manual annotation, each with inherent limitations. For example, Baysor [12] proposes to compare two segmentation methods by first identifying the consensus region segmented by both methods as well as the associated expression profile. Then, the correlation in gene expression between this consensus region and the areas excluded by each method is calculated. The method yielding higher correlation is considered superior. However, this approach cannot detect systematic errors shared by both methods, nor can it account for spatially heterogeneous gene expression within individual cells.

Another approach relies on simulations [14,25,8], which provide complete control over the ground truth. While simulations are useful, they may fail to replicate all the challenges encountered in experimental data. Finally, previous works [16] introduces quality metrics, such as shape and gene expression features, to identify outliers. However, these metrics serve more as a way to assess the plausibility of results, rather than offering a rigorous quantitative comparison against a ground truth.

We argue that the gold standard for validation of segmentation methods is the comparison to manually generated ground truth. To enable manual annotation of cell boundaries, even in regions of poor marker quality, we developed a novel visualization approach for RNA point clouds and incorporated it into our manual annotation workflow (Figure2A), as described below.

In order to visualize a large number of RNA species in the same image, we assign one color to each RNA species. However, assignment of random colors clearly cannot lead to an interpretable image. To overcome this limitation, we assign similar colors to genes that are co-expressed, based on the intuition that co-expressed genes are likely to belong to the same cell. By coloring each set of co-expressed genes differently, we can accurately distinguish cells having different gene co-expression patterns, as shown in Figure2B. Our novel co-expression visualization hence eases the annotation task on low quality staining area.

To implement this co-expression visualization, we first estimate a co-expression matrix using spatial correlation as a proxy for gene co-expression, as previously described [8] (see Methods5.6). Each gene is then represented by a co-expression vector, containing its spatial correlation values with other genes. These vectors are reduced to three dimensions using PCA, enabling their visualization as RGB colors (Figure2). Using this workflow, we annotated 724 cells from both high- and low-quality staining regions across five MERFISH datasets from different organs, breast, colon, melanoma and lung from VHFI, and the MERFISH Mouse Ileum dataset [12]. The number of annotated cells per datasets is summarized in Supplementary Table 2. The annotated datasets are available at: https://zenodo.org/records/14912364. The python code for RNA visualization is available at https://github.com/tdefa/gene coexpression visualization.

### 2.3 Benchmark on annotated datasets

Our annotated dataset provides a valuable resource for benchmarking segmentation methods. We compared RNA2seg with both image-based and point cloud-based segmentation approaches. For image-based methods, we evaluated the CNN-based Cellpose and the segmentation provided by Vizgen for the VHFI dataset. Of note, the Cellpose model used in this benchmark is also the Cellpose model used as a teacher to train RNA2seg. For point cloud-based methods, we tested Baysor [12] and our recently published method, ComSeg [8], both with nuclei as prior segmentation. Baysor groups RNA into single-cell expression profiles by identifying spatially homogeneous transcriptomic regions, while ComSeg leverages a spatial co-expression graph to assign RNAs to nuclei.

For staining-based methods, the segmentation output assigns each pixel to either a cell or the background. To evaluate performance, we calculated the best-matched Intersection over Union (IoU) between each annotated cell and the segmentation output. In the case of point cloud methods, pixels are not labeled and only RNAs are assigned to cells. In staining-based methods, segmentation assigns each pixel to either a cell or the background, whereas point-cloud methods associate RNAs with cells while leaving other pixels unlabeled. To ensure a fair comparison between staining-based and point cloud-based methods, we computed the best-matched RNA-level-IoU for each cell and reported the average across all cells. Likewise, pixel-level-IoU are reported for comparison of staining-based methods. We applied these methods to four VHFI datasets from human breast, colon, melanoma, and lung tissues (Figure 3 A-B, Supplementary Figure 1-2). While these datasets were part of the training set, the annotated regions were excluded during training, such that training and test sets were disjoint. On these datasets, RNA2seg-CNN and RNA2seg-Mamba demonstrated equal or superior performance compared to staining-based methods (Figure 3C). The performance advantage was particularly high on datasets where staining quality is often poor, like the VHFI Breast datasets (see Supplementary Figure 3). In contrast, RNA2seg, Cellpose and the segmentation published with the VHFI performed similarly on the VHFI lung dataset, where staining quality is high. This aligns with our ablation study (Supplementary Figure 4), which shows that the performance gap between RNA2seg models with and without RNA point cloud is small on the VHFI lung dataset, whereas it is high on the VHFI breast dataset, where heterogeneous staining quality makes the additional RNA positional information valuable.

**Fig. 3.**
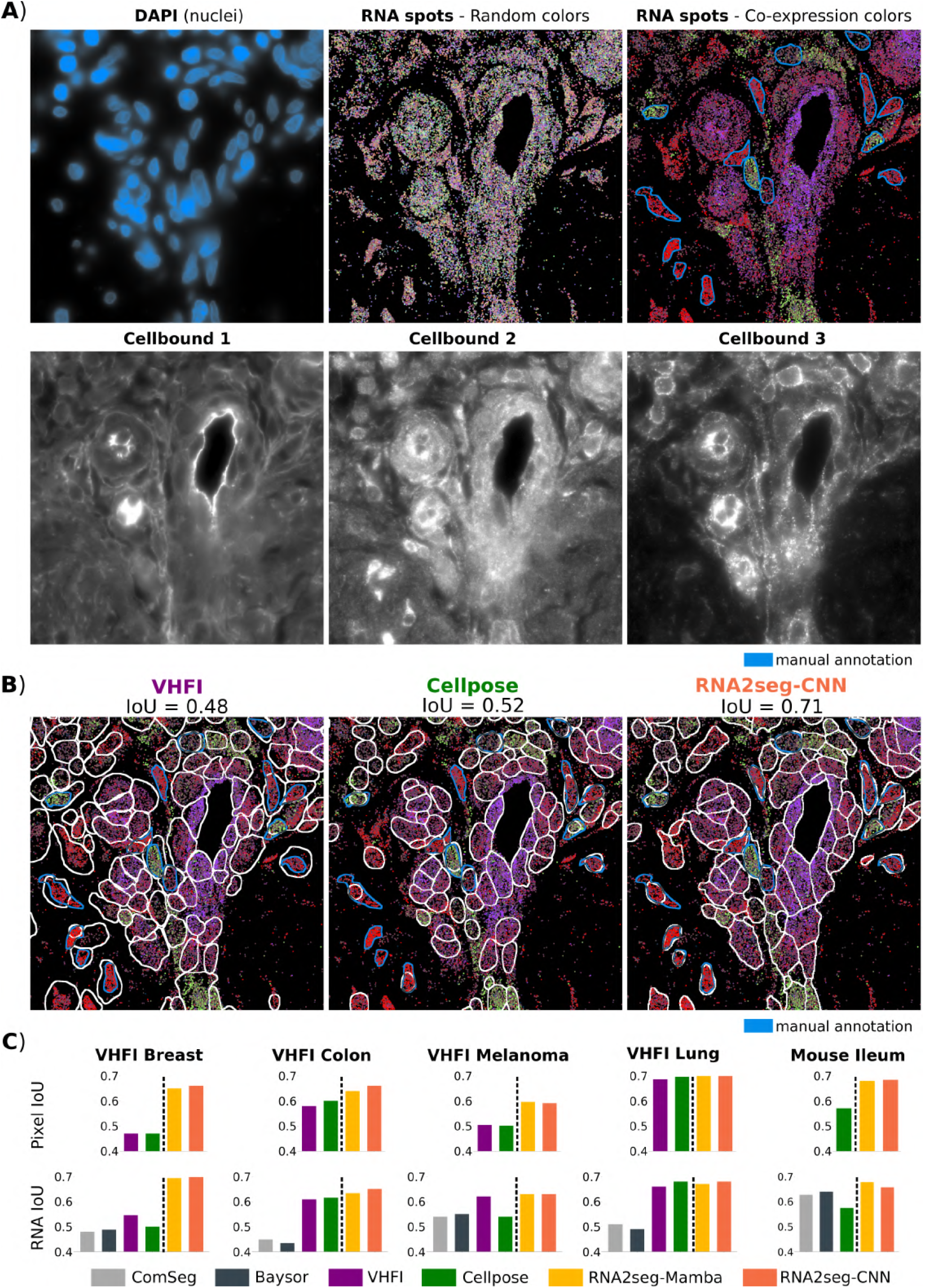
Benchmark of different IST cell segmentation methods. **A)** Example of stainings and RNA point clouds. From left to right in the first row: DAPI staining, RNA spots with random color, RNA spots with co-expression color, and manual annotation in blue. The second row displays the three different available stainings in the VHFI dataset. **B)** Example of segmentation from VHFI, Cellpose, and RNA2seg-CNN **C)** Benchmark of Mean Pixel IoU and RNA IoU across different methods and datasets.

**Fig. 4.**
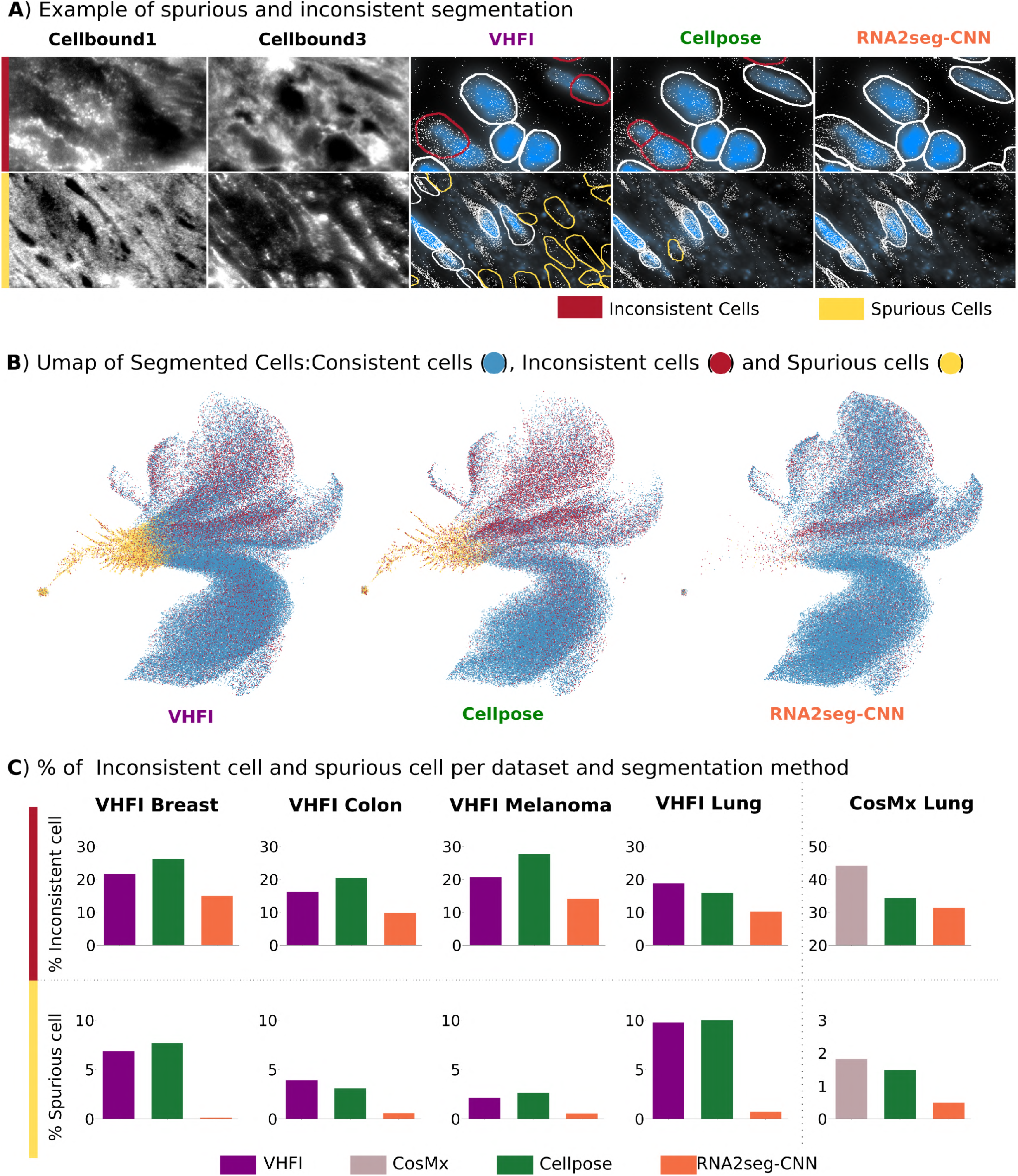
Spurious and inconsistent cells. **A)** Example of spurious and inconsistent segmentation. **B)** Umap of segmented cells: consistent cells are colored in blue, inconsistent cells in red, and spurious cells in orange. **C)** % of inconsistent and spurious cells for each dataset and segmentation method.

Point cloud-based methods, Baysor and ComSeg, performed worse than image-based methods on datasets with high-quality staining, such as lung tissue. However, their performances are only slightly inferior on datasets with challenging staining, like breast and melanoma.

We also annotated cells in the mouse Ileum dataset [12], which was not part of the training set and features a different staining scheme. Unlike the training datasets, it includes only one cell boundary and a DAPI staining, but no PolyT staining. Despite these differences, RNA2seg outperformed ComSeg, as well as the Baysor and Cellpose segmentations published in the original study. Of note, the Cellpose segmentation used in this benchmark comes from the Cellpose model specifically fine-tuned on the Mouse Ileum dataset by the original authors. These results highlight the generalization capabilities of RNA2seg, demonstrating its effectiveness on an unseen dataset from a different species, a different tissue type, and with different staining conditions.

We also compared different deep learning architectures. Overall, RNA2seg-CNN and RNA2seg-Mamba demonstrate comparable performance. However, RNA2seg-Mamba is significantly more computationally demanding, with seven times more parameters than RNA2seg-CNN (6.6 million vs. 44.2 million). Therefore, for the remainder of this study, we focus exclusively on RNA2seg-CNN, referred to as RNA2seg. Of note, the performance of RNA2seg is stable across different training initializations, as shown in Supplementary Figure 5. The performance of RNA2seg-CNN displayed in Figure 3 are the average score from of 10 independently trained models. All the segmentation generated for this benchmark are available at: https://zenodo.org/records/14916899

**Fig. 5.**
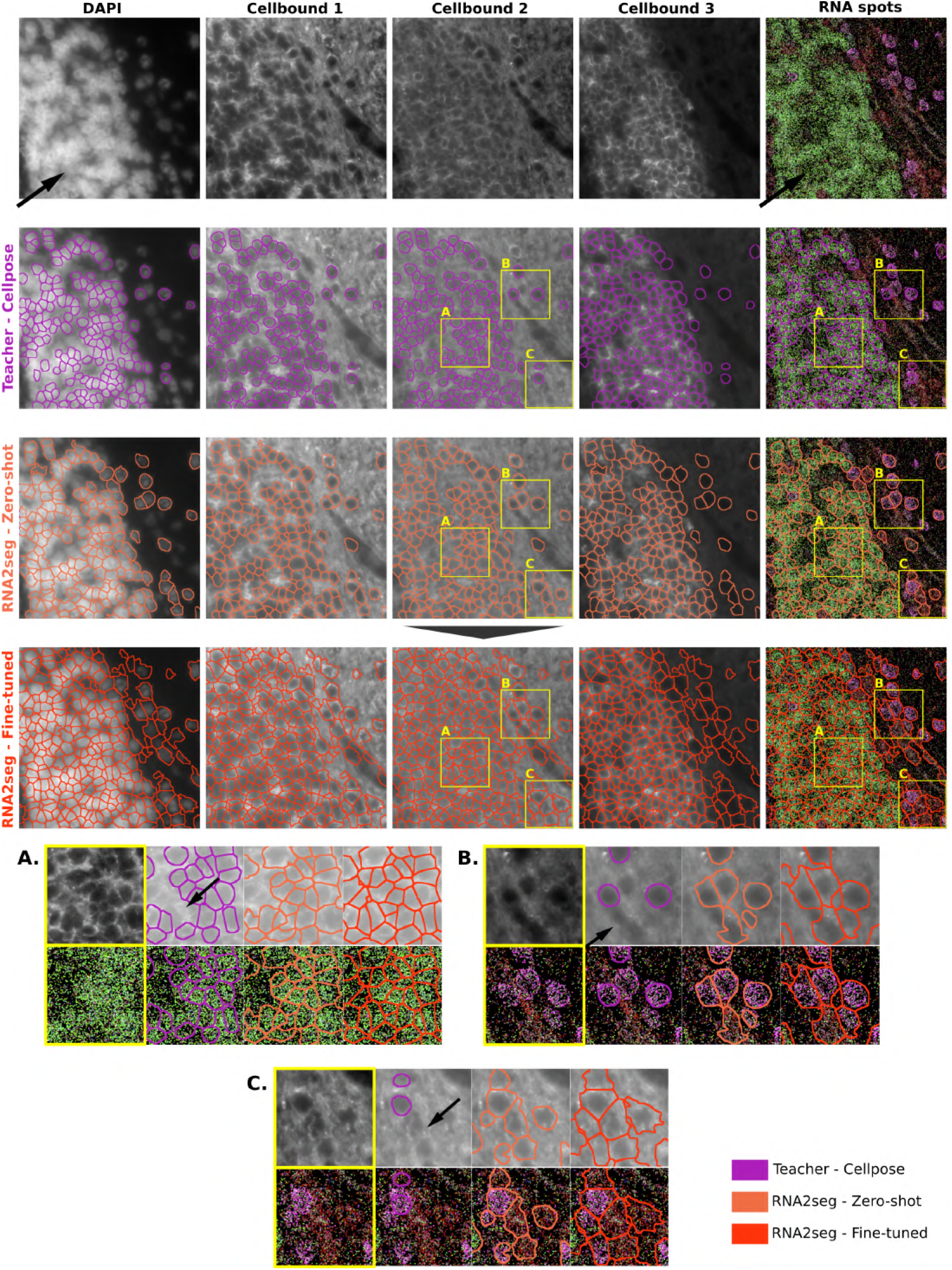
Visualization of RNA2seg segmentation using zero-shot and few-shot-learning for fine-tuning on the Hamster Brain dataset. The first row presents the input data: DAPI, three different cell boundary stainings, and RNA spots colored using our co-expression visualization. The second row displays the Cellpose teacher segmentation used for fine-tuning RNA2seg. It then shows in the third row the RNA2seg zero-shot segmentation results. The fourth row shows the segmentation done by our fine-tuned RNA2seg using the Cellpose teacher segmentation of the second row. The figure includes three zoomed-in crops (**A, B, C**) extracted from the initial data, showcasing the three segmentation results (teacher segmentation, zero-shot RNA2seg, and fine-tuned RNA2seg) side by side. This visualization highlights the significant improvement achieved through few-shot-learning, enhancing both the alignment with cell boundary stainings and RNA distribution. Specifically, plot **A** highlights a region where both Zero-Shot RNA2seg and the teacher model miss cells, while fine-tuned RNA2seg successfully segments them. Plots **B** and **C** demonstrate improved boundary placement in areas with weak staining, even where the teacher model fails. Overall, this figure illustrates that few-shot-learning remains effective even when the teacher segmentation overlooks cells, particularly in poorly stained regions. Fine-tuning enables the model to adapt efficiently to a new dataset while maintaining good performance.

### 2.4 Comparison of different RNA image representations

In the above results, the RNA representation used by the network exclusively relates to RNA position and disregards RNA species identity. However, RNA identity can be integrated into the model along with positional information by assigning a unique color to each RNA species. In order to test whether this is beneficial to the overall algorithm performance, we used first the encoding initially developed for visualization (Section 2.2), where we assign similar colors to RNA species that are co-expressed in the tissue. Secondly, we evaluated an approach where the colors assigned to RNA species are treated as learnable parameters during training. The goal of this approach is to let the network determine the optimal RNA species colors for segmentation.

While the inclusion of co-expression information in RNA visualizations can aid manual annotation by humans, our ablation study (Supplementary Figure 4) demonstrates that it does not improve RNA2seg’s performance. We obtain a comparable performance for RNA2seg with positional-only data, co-expression-based coloring, and learned RNA species colors. This suggests that the RNA2seg already captures the usefull RNA identity information inherently from RNA positional data.

Additionally, a potential limitation of using RNA colors to encode RNA species identity instead of solely using RNA positions is that RNA2seg may not generalize well to new datasets. Indeed, different datasets might spatially resolve different RNA species, leading to novel color patterns that could require re-training. Similarly, co-expression-based RNA visualizations degrade model performance in zero-shot settings, as demonstrated by the results on the mouse ileum dataset (Supplementary Figure 4), which was not part of the training data. This performance decline occurs because co-expression visualizations rely on dataset-specific gene panels, leading to different color patterns across datasets with different panels. Such variability hinders the model’s ability to generalize to unseen data. Finally, this comparison of various RNA image representations confirms that relying solely on RNA positional information is a relevant approach for RNA2seg.

### 2.5 Quantification of cell-level segmentation errors

So far, our evaluation metrics measured inaccuracies in contours or RNA assignment. Next, we turned to cell-level metrics providing further quantitative comparisons between the benchmarked methods.

1. **Spurious cells** are defined as cells that do not intersect with any nucleus and contain fewer than a minimal number of RNA molecules (see Methods 5.8). Such cells are likely false-positive segmentations.
2. **Inconsistent cells** are defined as cells that intersect with a nucleus but fail to fully enclose it and cells that contain several nuclei (see Methods 5.8). Such cases are likely indicative of segmentation inaccuracies.

Examples of spurious and inconsistent cells are illustrated in Figure 4A. A UMAP visualization of the segmented cells from each method, highlighting spurious and inconsistent cells, is presented in Figure 4B. We observe that inconsistent cells and in particular, spurious cells show a very specific gene expression pattern, suggesting that their identification and exclusion from the analysis are essential for downstream analysis.

Both metrics are based on prior nucleus segmentation derived from DAPI staining, which is typically more stable and reliable across tissues and cell types compared to cell boundary staining. This DAPI staining stability allows us to assume that nuclei segmentation is accurate. Under this assumption, we calculate the proportion of spurious and inconsistent cells as a surrogate measure of segmentation quality.

Across the four datasets analyzed, RNA2seg demonstrates an almost negligible percentage of spurious cells (Figure 4C). While spurious cells can be eliminated through post-processing, their low occurrence indicates that RNA2seg effectively associates RNA density variations with cell instances, achieving one of the method’s key objectives. Furthermore, RNA2seg exhibits a lower percentage of inconsistent cells compared to other methods, meaning it produces biologically coherent segmentations with fewer implausible segmentation results (Figure 4C). This highlights the model’s ability to generate more accurate and meaningful segmentations, which is essential for downstream analyses.

### 2.6 Automatic fine-tuning of RNA2seg

RNA2seg was trained on publicly available datasets. However, new datasets can display cell diversity and staining variability that differ from these training data. In this section, we apply RNA2seg to out-of-training-distribution data, using an in-house Hamster MERFISH dataset as an example. This dataset displays significant differences from the training data, with locally reduced RNA density and variable DAPI signal quality. Further, the nuclei in certain regions are densely packed and indistinguishable from one another (Figure 5, first row, arrows).

We first applied Cellpose on the cell boundary staining 1, which is one of the proposed standard workflows by the vendor (Figure 5, second row). Although Cellpose effectively delineates many cells, some remain unsegmented (see Figure 5, detail A-B-C, arrows). We then applied RNA2seg without retraining in a zero-shot setting (Figure 5, third row). While cell segmentation was generally accurate, not all cells are segmented, especially in areas where the data deviates significantly from the training distribution. This issue is particularly noticeable in regions where the DAPI signal is near saturation and the RNA density is low.

We then explored whether few-shot learning could increase the usability of our model for such unseen data. In short, this strategy allows the model to be fine-tuned with minimal training and low computational demands, requiring only a few iterations on new targeted data to quickly adapt to an out-of-distribution dataset. However, even such minimal retraining requires annotated data. To remove the need to manually curate such data, we leveraged our automatic student-teacher training approach to fine-tune RNA2seg on our Hamster MERFISH dataset by using the Cellpose segmentation (see Method section 5.9). Importantly, this annotation is not always accurate and incomplete but can be obtained fully automatically. After retraining for 10 minutes on our P100 GPU (see Method section 5.9), RNA2seg yielded significantly improved results, especially in regions that were previously poorly segmented (Figure 5, fourth row).

In summary, the results presented in Figure 5 show that automatic fine-tuning of RNA2seg leads to better segmentation results on our in-house Hamster Brain dataset. Surprisingly, Figure 5 also shows that even though the Cellpose teacher model sometimes misses cells and struggles to segment areas with weak staining, the fine-tuned RNA2seg student model still produces consistent segmentation results in these areas (Figure 5, fourth row). This supports the efficiency of our student-teacher training approach and indicates that our automatic fine-tuning allows the model to easily adapt to new tissue types.

## 3 Discussion

Cell segmentation is currently a major bottleneck in using IST data for spatial biology. Segmentation errors can propagate through downstream analyses, leading to inaccurate assessments of subcellular RNA localization, erroneous cell-level gene expression measurements, and ultimately incorrect cell type classification.

Here, we present RNA2seg, which, to the best of our knowledge, is the first cell segmentation method for IST data that integrates DAPI staining, multiple cell membrane stainings, and the spatial distribution of RNAs. By leveraging these complementary data sources, RNA2seg is particularly effective in regions with low staining quality, where the last generation of cell segmentation methods for fluorescence microscopy data, such as Cellpose, are bound to fail. At the same time, RNA2seg fully utilizes high-quality staining data where it is available.

RNA2seg relies on a teacher-student framework and can therefore be trained without relying on manual annotation. This is especially important for IST data, where manual annotation is exceptionally time-consuming. Furthermore, large deep learning models achieve their full potential only when trained on extensive annotated datasets. Our framework allows RNA2seg to be trained on more than 4 million cells across diverse tissues, gene panels, and membrane markers, ensuring broad representation of the expected input data distribution and significantly improving robustness.

Validation of segmentation algorithms for IST data is complicated, as manual annotation is particularly difficult, requiring interpretation of point clouds, which is notoriously difficult, and integrating information scattered over several staining channels. Most authors have therefore opted for indirect measurements to assess segmentation accuracy. We argue that the best way of assessing the quality of segmentation methods is still to compare them to a manually generated ground truth. For this reason, we developed a new visualization scheme, where RNAs are colored according to their co-expression. This allows to utilize RNA point clouds in an efficient way for identifying cellular boundaries, in particular in regions of poor staining quality. We have annotated 724 cells across different tissue types and experimental setups, which we make freely available to the scientific community (https://zenodo.org/records/14912364). We expect that this manually curated dataset will ease future developments and in particular serve as a basis for benchmarks in the field, which are today still difficult to achieve.

We demonstrate that incorporating RNA species information into our algorithm did not improve its performance. This result is unexpected, since it could be assumed that gene expression data would provide valuable information for segmenting cells with distinct transcriptional profiles. Several potential reasons could explain this observation. First, RNA density alone may be already a sufficiently informative feature, making the additional inclusion of RNA species data redundant. Indeed, improvements are only expected in scenarios where neighboring cell types cannot be adequately segmented using membrane and RNA positional data alone — likely representing a small subset of the observed segmentation errors. Another explanation is that the current algorithmic strategies for leveraging RNA species data may still be suboptimal.

In a series of validation experiments, we showed that RNA2seg outperforms state-of-the-art methods for cell segmentation in IST data, both on data from the same experiments but withheld during training and in a zero-shot-learning setting on IST data of a different organism, a different set of membrane markers and a different gene panel.

RNA2seg was designed as a flexible and generalist model, capable of adapting to a wide variety of datasets. First, it can accommodate an arbitrary number of input channels, enabling it to fully leverage all available data. Second, its RNA representation is independent of the specific gene panel, ensuring that the model remains applicable across datasets with varying gene compositions. Third, RNA2seg demonstrates robust performance in a zero-shot learning setting. Finally, to further tailor the model to user-specific datasets, we proposed a few-shot learning approach that is fast, highly effective, and requires no manual annotations. Together, these features make RNA2seg a powerful and versatile tool for cell segmentation across diverse biological sample types.

While RNA2seg demonstrates competitive performance in cell segmentation, it has several limitations. First, it is currently restricted to 2D data. Although many commercial systems currently perform 2D measurements or probe only thin 3D volumes, the ability to analyze thick 3D samples will likely become increasingly important, particularly for studying intact organoids or tumoroids [26]. Second, we have chosen to represent RNA point clouds as images. When using large marker panels, intra-cellular RNA point clouds are typically dense rather than sparse, making this representation a logical choice. However, exploring architectures that directly process point clouds could be a promising direction, as we have previously proposed for classifying subcellular localization patterns [27]. While integrating such an approach with the CNN-based architecture used for membrane stainings presents conceptual challenges, it could potentially enhance efficiency. Finally, while we did not find that a recent Mamba-based architecture provided any tangible advantage in segmentation performance, leading us to opt for a more parameter-efficient CNN-based architecture, this choice may be reconsidered with the emergence of foundation models for segmentation [28]. These models, predominantly based on transformer architectures, could further enhance the segmentation of IST data, potentially shifting the balance away from CNNs.

## 4 Conclusion

In this study, we introduce RNA2seg, a generalist segmentation algorithm designed for image-based spatial transcriptomics. Unlike existing methods, RNA2seg integrates information from multiple staining and RNA localization to achieve accurate cell segmentation. To train RNA2seg without manual annotations, we developed a student-teacher training scheme, enabling the model to learn from a large and diverse dataset. This approach ensures that RNA2seg is generalizable and capable of performing zero-shot segmentation on unseen datasets. We argue that manual annotations remain essential for benchmarking, even though they are notoriously challenging to generate for IST data. To address this, we designed a novel visualization scheme, which enabled us to annotate more than 700 cells across diverse IST datasets — providing a valuable resource for future benchmarking studies. Finally, RNA2seg outperforms existing methods on various segmentation metrics, highlighting its utility in this fundamental task for downstream single-cell analysis. In practice, RNA2seg is implemented for OME-zarr to ease large scale applications and is available as a pip package: https://github.com/fish-quant/rna2seg

## 5 Methods

### 5.1 Method overview

RNA2seg is a deep learning-based cell segmentation method that takes as input DAPI channel *X*_*DAPI*_, RNA point clouds 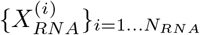, and optionally additional membrane staining channels (MSC) 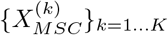. The output is a cell segmentation mask.

A core element of our approach is an efficient teacher-student framework, which enables the training of a dedicated segmentation method without requiring manual annotation.

Cell segmentation from fluorescent membrane staining channels is a well-studied problem, where previously published and trained convolutional neural networks (CNNs) have demonstrated performance close to human-level accuracy [9, 10]. However, the segmentation quality of such methods is conditioned on the staining quality which tends to be highly heterogeneous in IST data. Consequently, segmentation results can be excellent in some locations and very poor in others.

To generate a high quality training set automatically in a teacher-student framework, our strategy leverages Cellpose to generate an initial segmentation of nuclei and cells from DAPI and MSC. We then apply a series of rules to filter out segmentation results likely to be incorrect or inaccurate. Using the filtered training labels, RNA2seg learns the RNA point cloud density patterns corresponding to cells. Once trained, RNA2seg performs cell segmentation even in areas lacking high-quality membrane staining.

### 5.2 Teacher-network: generation of the training set

In order to generate the labeled data for training RNA2seg, we first apply teacher pre-trained networks for cell and nucleus segmentation and then select among the segmented cells high-confidence objects (*C*_*v*_), as described below. Additionally, we identify high-confidence background areas (*B*_*v*_). Then, inspired by SketchPose [24], we generate partial back-propagation masks to train the model exclusively on high-confidence cells (*C*_*v*_) and background areas (*B*_*v*_).

#### 5.2.1 Candidate cell segmentation with a pre-trained network

We use the Cellpose cyto3 [10] pre-trained model for segmenting cells. Each MSC is segmented individually, using a two-channel image formed by concatenating *X*_*DAPI*_ and 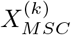. For DAPI segmentation, we also utilize the Cellpose nuclei [10] pre-trained model.

#### 5.2.2 Background segmentation

We use a training scheme for partial annotation, i.e. pixels are not systematically negative if they are outside high-confidence cells. To identify the background regions (*B*_*v*_) devoid of cells, we rely on RNA density as an indicator. The underlying assumption is that areas with low RNA density correspond to background regions without cells. To estimate RNA density for each pixel, we apply a Gaussian kernel density estimation with a standard deviation of *σ* = 10 pixels. Background regions are then defined as the set of pixels with RNA density below a threshold *T*_*b*_. This threshold *T*_*b*_ is defined as the 5%-percentile of RNA densities inside the nucleus.

#### 5.2.3 Selection of consistent cell segmentations

In this section, we describe how we compute the set of high-confidence cells, denoted as *C*_*v*_, from the cell segmentation provided by the teacher network. Our objective is to exclude poorly segmented cells, particularly those from regions with weak cell boundary staining. Our approach assumes that the nucleus segmentation is accurate, which is often the case, as the DAPI channels are usually of good quality. Valid cells are identified based on the consistency between cytoplasm and nucleus segmentation and the RNA density. The agreement between a segmented cell and an independently generated nucleus segmentation thus serves as a surrogate measure of correctness.

To categorize cell-nucleus segmentation consistency, we first define relationships between cells and nuclei based on area overlap thresholds:

- A cell is considered to **contain a nucleus** if at least *T*_*contain*_ = 95% of the nucleus area overlaps with the cell.
- A cell is considered to **intersect a nucleus** if more than *T*_*intersect*_ = 1 − *T*_*contain*_% of the nucleus area overlaps with the cell.

Among the cells segmented by the teacher network, the following cells are retained:

1. Cells that contain one and only one nucleus.
2. Cells that do not intersect with a nucleus.
3. Nuclei that do not intersect with more than one cell.

Each rule addresses a specific segmentation scenario:

1. The first rule ensures that cells with consistent nuclear and cytoplasmic segmentation are retained.
2. The second rule includes cells without an associated nucleus to account for cases where the nucleus is missing from the tissue section.
3. The third rule addresses regions with poor MSC quality, which may result in cellular regions missed by the teacher network. In such cases, the nuclear regions are used instead. Including nuclear regions in the training labels does not compromise the ground truth, thanks to our partial back-propagation method described below. We assume however that cell and nucleus center are close to each other.

Finally, we exclude from *C*_*v*_ any cell without nucleus or nucleus without cell with more than *T*_*bg*_ = 5% overlap with background regions, as such overlap indicates potential segmentation errors.

#### 5.2.4 Generated training label in this study

We segment cells from different MSC independently and generate training targets for each segmentation result separately by applying the rules described above. The resulting training labels are then merged into a comprehensive training dataset. Since different MSC can be complementary and no single marker consistently provides superior results across all regions, incorporating training labels from multiple MSC ensures a more extensive and robust training dataset.

For the MERFISH datasets, we generate the training target from the cell boundary staining 1 and 3. For the cosmx dataset we generate the training target from all of the three available immunostainings. The training labels used in this study are available at: https://zenodo.org/records/14916899

### 5.3 RNA image generation

We transform RNA point cloud data into image data by assigning to each pixel the number *n* of RNA molecules it contains, irrespective of their species. Following the approach described in [15], we then apply a max filter with *k* = 2 and Gaussian kernel filter with *σ* = 2 to smooth the data. The output is an 1 × *H* × *W* image which we denote *I*_*RNA*_.

### 5.4 RNA2seg network and loss

The RNA2seg network takes as input two images: a staining image *X*_*S*_ including all MSC and the DAPI channel of size (*K* + 1) × *H* × *W*, where *K* corresponds to the number of MSC, and an RNA image *I*_*RNA*_ of size 1 × *H* × *W*.

To allow RNA2seg to handle an arbitrary number of stainings, *X*_*S*_ is first processed using a ChannelNet architecture [20], which outputs a fixed-size image *I*_*S*_ of size 3 × *H* × *W*, independent of the number *K* of MSC.

*I*_*S*_ is then concatenated with *I*_*RNA*_ and passed into a U-Net backbone, which can either be a Mamba-U-Net or a CNN-based U-Net architecture adapted from the Cellpose U-Net.

As output, the model produces an image of size 3 × *H* × *W*, where the first channel indicates the probability that a pixel belongs to a cell, and the next two channels provide the horizontal and vertical gradients pointing to the cell center, as implemented in Cellpose [10].

The model’s loss function consists of two components: a cross-entropy loss on the cell probability output and a mean squared error (MSE) loss on the predicted gradient that points toward the cell centers.

### 5.5 RNA2seg training

To enable RNA2seg to effectively segments areas with low-quality or missing cell boundary staining, we applied dropout to the channels of *I*_*S*_ during training. Specifically, membrane staining image channel were replaced by empty images with a dropout probability of 0.25, which effectively emulates missing membrane staining signals.

Additionally, we applied random brightness-contrast adjustments, Gaussian blur filtering, random rotations, an random crop masking of the staining images to enhance robustness.

RNA2seg was trained on 64k partially labeled images of size 1200 × 1200, resized to 512 × 512. The training was conducted with a batch size of 8, using the Adam optimizer [29] with a learning rate of 0.001. We reserved 5% of the training dataset for validation and selected the best model based on validation performance after 30 epochs.

### 5.6 Co-expression Visualization

To validate RNA2seg, we manually annotated cells by simultaneous inspection of all channels (DAPI, MSC and the generated RNA density channel). However, as shown in Figure2B, accurately annotating cell boundaries solely with staining can be challenging, in particular in regions of low staining quality. To facilitate the annotation process, we assign similar colors to RNAs from co-expressed genes, based on the assumption that RNAs from genes co-expressed are likely to belong to the same cell.

First, we estimate co-expression using the spatial arrangements of detected RNA molecules in the image as described in our previous work [8]. In brief, by analyzing the spatial distances of the detected RNA positions, we can estimate a spatial co-expression value for each gene pair that recapitulates how likely transcripts of the two genes appear in close proximity. This leads to a co-expression matrix 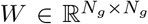 where *N*_*g*_ is the the number of genes in the IST dataset and the *w*_*i,j*_ correspond to the spatial co-expression of gene *i* and *j*. Each gene *i* is characterized by its coexpression profile (columns *W*_*i*_ of the matrix *W*):

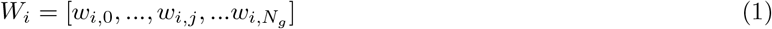

Application of Principal Components Analysis (PCA) to *W* allows us to map each column vector *W*_*i*_ to a 3-dimensional vector that is then used as an RGB color code of the corresponding RNAs.

### 5.7 RNA2seg with colors for each RNA species color

We also first explored whether this Co-expression visualization scheme could be beneficial for representing RNA distributions during training. Hence, instead of using a one-channel image to encode the RNA positions, as described above, we represented each RNA species as a 3-dimensional vector, interpretable as an RGB color. Our hypothesis was that in addition to RNA density, the differences in transcriptional programs between cells of different type should in principle bear useful information for segmentation.

Alternatively, we tested to treated RGB vectors of each RNA species as learnable parameters that are optimized during training.

To implement these two solutions, for each RNA position, its corresponding 3-dimensional vector is incorporated into the RNA image *I*_*RNA*_, with dimensions 3 × *H* × *W*. The segmentation network’s first convolutional layer is modified to accommodate the dimensions of *I*_*RNA*_. If multiple RNA species occupy the same position in the image, one is randomly selected for representation.

### 5.8 Validation and metrics

#### 5.8.1 Intersection over Union (IoU)

Our annotations are done on image of 1200×1200 pixels and are partial as only non ambiguous cell were annotated. We computed the intersection over union (IoU) for each annotated cell and its best matched segmented cell. The best match is calculated by finding the cell that maximizes the IoU at the pixel (pixel IoU) or the RNA level (RNA IoU).

#### 5.8.2 Spurious and inconsistent cells

As an additional validation metric, we propose to calculate the number of spurious and inconsistent cells:

- **Spurious cells** are defined as segmented cells that do not intersect a nucleus and have an RNA count below *T*_*spurious*_, which we set to the 5-percentile of RNAs inside nuclei. Spurious cells likely correspond to false positives.
- **Inconsistent cells** are defined as segmented cells that intersect at least one nucleus without fully enclosing it or segmented cells containing more than one nucleus, with respect to the defined thresholds *T*_*contain*_, *T*_*intersect*_.

### 5.9 Automatic fine-thuning on our in-house Hamster MERFISH dataset

We first applied the Cellpose DAPI model to segment nuclei and the Cellpose cyto3 model to delineate cell boundaries from the Hamster MERFISH dataset [10]. These segmentations were used to generate a training dataset of 742 partially labeled images following the method described in Section 5.2. We then fine-tuned our pre-trained RNA2seg-CNN over 20 epochs, a process that took less than 10 minutes on our P100 GPU.

### 5.10 Benchmarked method hyper-parameters

For the VHFI dataset and the lung CosMX, we apply a Cellpose cyto3 with a diameter parameter of 60 on each available cell boundary staining. We kept the segmentation from the cell boundary staining performing the best. We apply Baysor with the nuclei segmentation as input and a confidence parameter of 1. We apply ComSeg with mean cell diameter of 7µm.

### 5.11 Datasets

For the training, We use seven dataset from the Vizgen’s Human FFPE Immuno-oncology Data Release (VHFI) and a Human Lung CosMx dataset from [11] detail in the Supplementary Table 1. For the Human Lung CosMx dataset cell, we retain a subset of the dataset named “Lung5 Rep1”.

**Table 1.** Summary of the datasets used for the training. The number of cells of each VHFI dataset comes from the Vizgen data released. The number of cells for the CosMx dataset is inferred with Cellpose on membrane staining.

| Dataset name | Imaging approach | number of Cells |
| --- | --- | --- |
| VHFI Breast | MERFISH | 713,121 |
| VHFI Lung | MERFISH | 353,762 |
| VHFI Col on | MERFISH | 677,451 |
| VHFI Melanoma | MERFISH | 468,138 |
| VHFI Ovarian | MERFISH | 358,485 |
| VHFI Prostate | MERFISH | 721,668 |
| VHFI Uterine | MERFISH | 843,285 |
| CosMx-lung (Lung5_rep1) [1] | CosMx | 87955 |

**Table 2.**
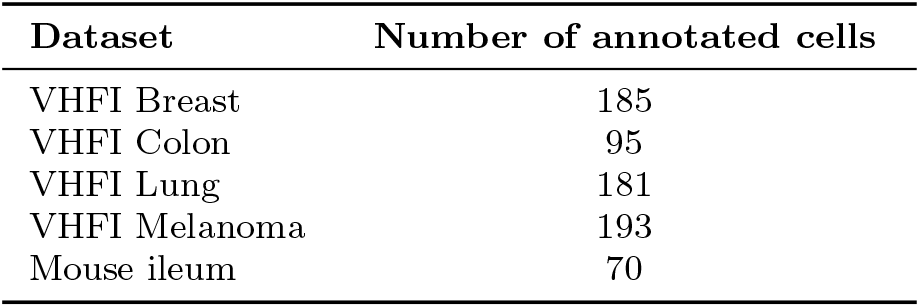
Numbers of manually annotated cells per dataset.

## 6 Data availability

RNA2seg is available as a fully documented Python pip package (https://github.com/fish-quant/rna2seg) and the weights of the different trained RNA2seg networks can be also found here: https://huggingface.co/aliceblondel/RNA2seg.

Our manually annotated dataset of cells is available at https://zenodo.org/records/14912364.

The Python code of our novel visualization is available at: https://github.com/tdefa/gene coexpression visualization The segmentation results computed for the benchmark are available at: https://zenodo.org/records/14916899

## 7 Acknowlegement

This work has received financial support through the Agence Nationale de la Recherche (ANR) for grants (LUSTRA, reference ANR-19-CE14-0015-04 to J.-A.L.-V., C.F., F.Mu., and T.W.) and (TRANSFACT, reference ANR-19-CE12-0007-02, to F.Mu. and T.W.). F.Mu. and C.W. acknowledge funding by Institut Pasteur. Furthermore, this work was supported by the French government under the management of Agence Nationale de la Recherche as part of the “Investissements d’avenir” program, reference ANR-19-P3IA-0001 (PRAIRIE 3IA Institute). Furthermore, this work was supported by a government grant managed by the Agence Nationale de la Recherche under the France 2030 program, with the reference number ANR-24-EXCI-0005. The authors also acknowledge access to the HPC resources of IDRIS under the allocation AD011015678 made by GENCI. G.D.M. acknowledges funding from the Fondation pour la Recherche Médicale (grant ANRS MIE202112015304).

## Supplementary information

### 1 Supplementary

**Fig. 1.**
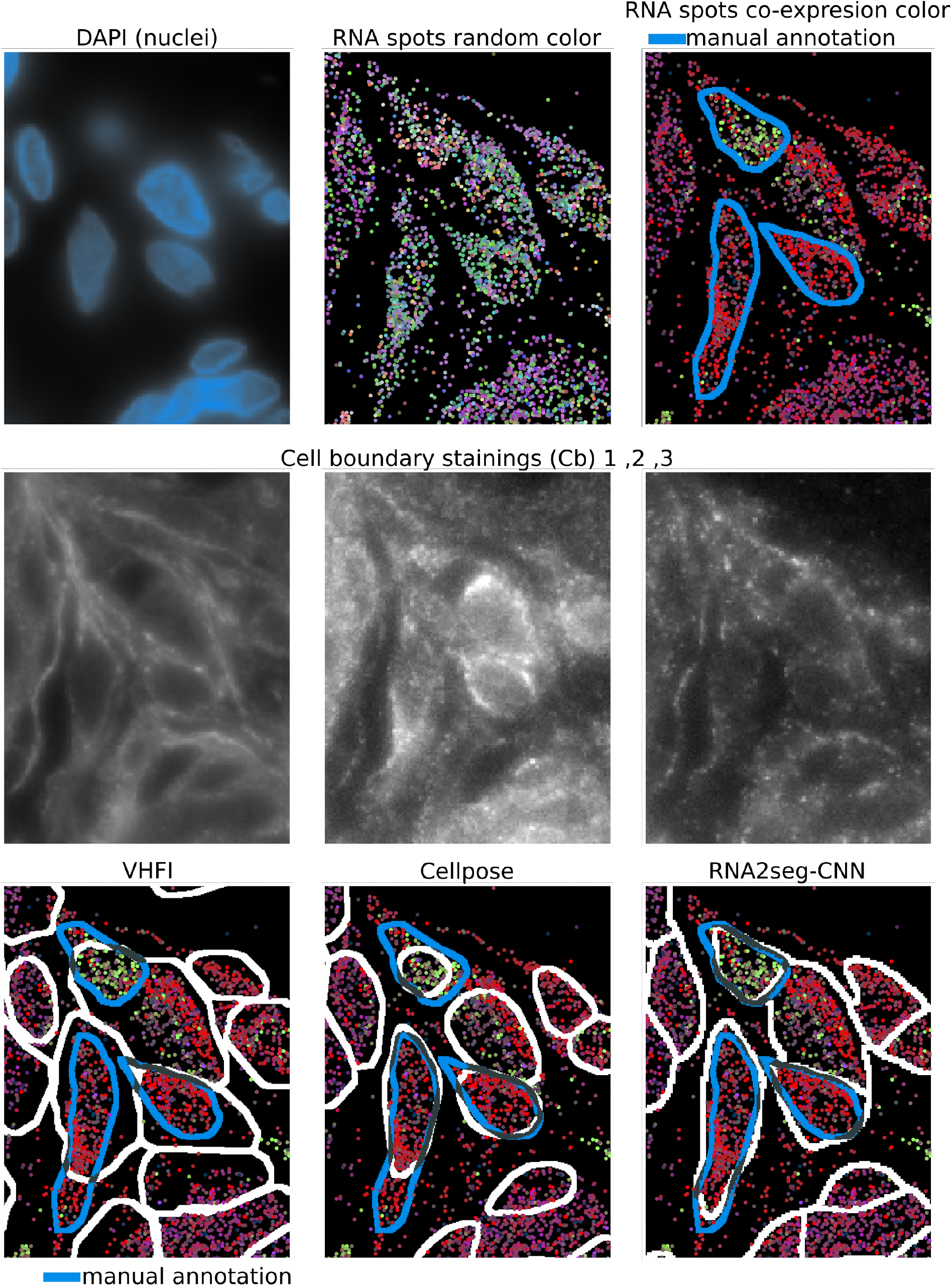
A) Example of staining and RNA point cloud. From left to right in the first row: DAPI staining, RNA spots with random color, RNA spots with co-expression color and manual annotation in blue. The second row display the three different available stainings in the VHFI dataset. The third row display examples of segmentation from VHFI, Cellpose, and RNA2seg-CNN

**Fig. 2.**
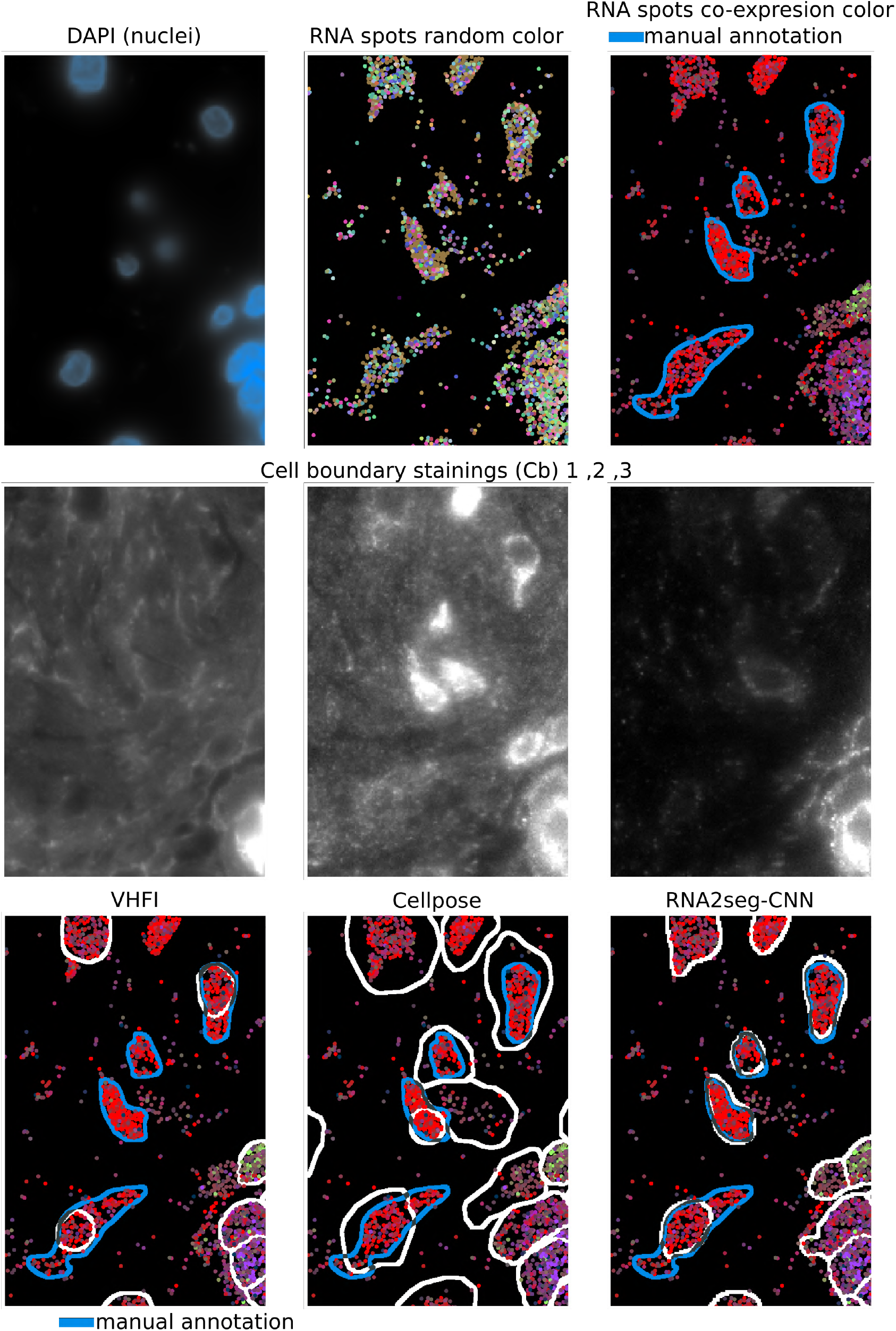
A) Example of staining and RNA point cloud. From left to right in the first row: DAPI staining, RNA spots with random color, RNA spots with co-expression color and manual annotation in blue. The second row display the three different available stainings in the VHFI dataset. The third row display examples of segmentation from VHFI, Cellpose, and RNA2seg-CNN

**Fig. 3.**
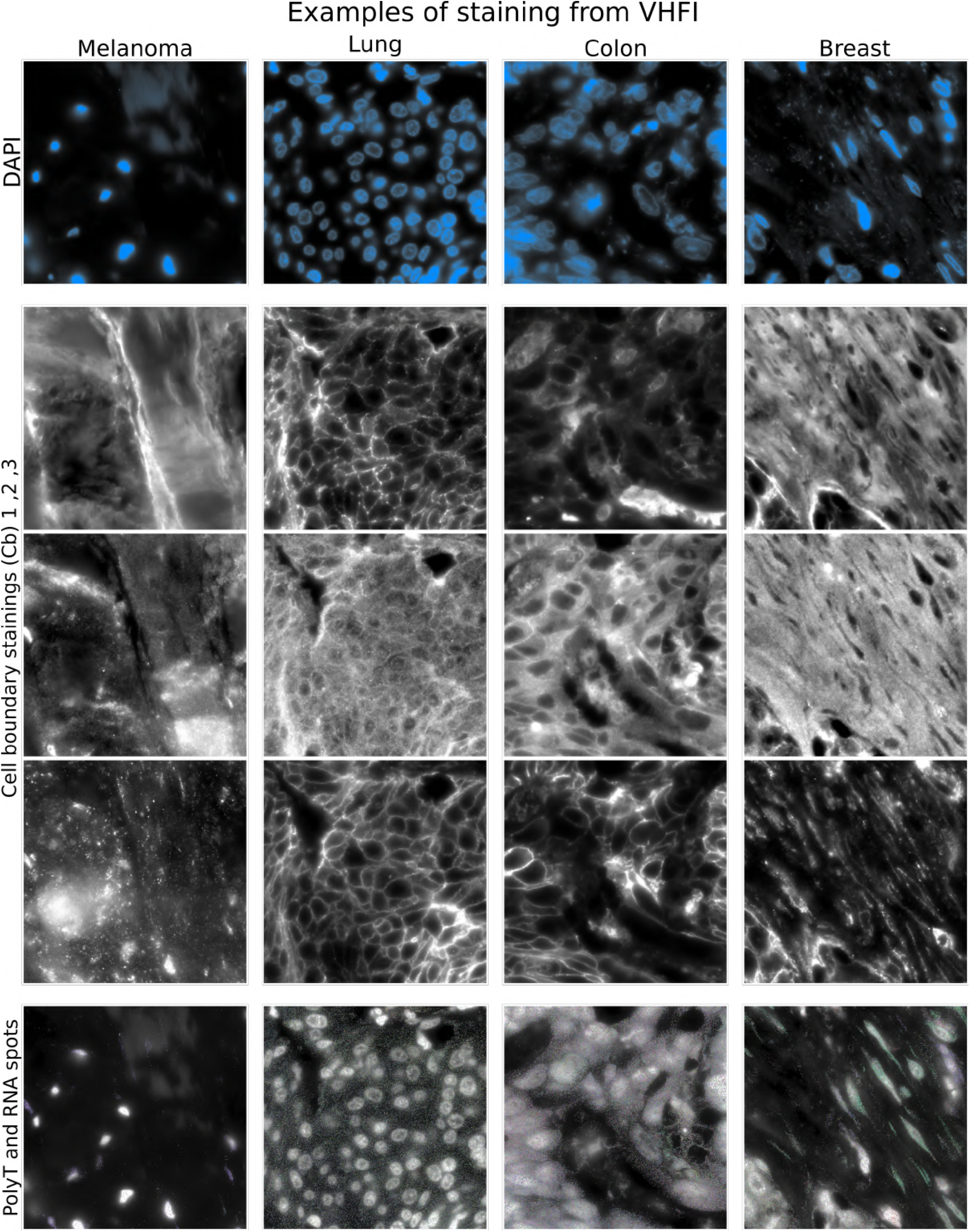
Examples of staining and detected RNA spots from the VHFI datasets.

**Fig. 4.**
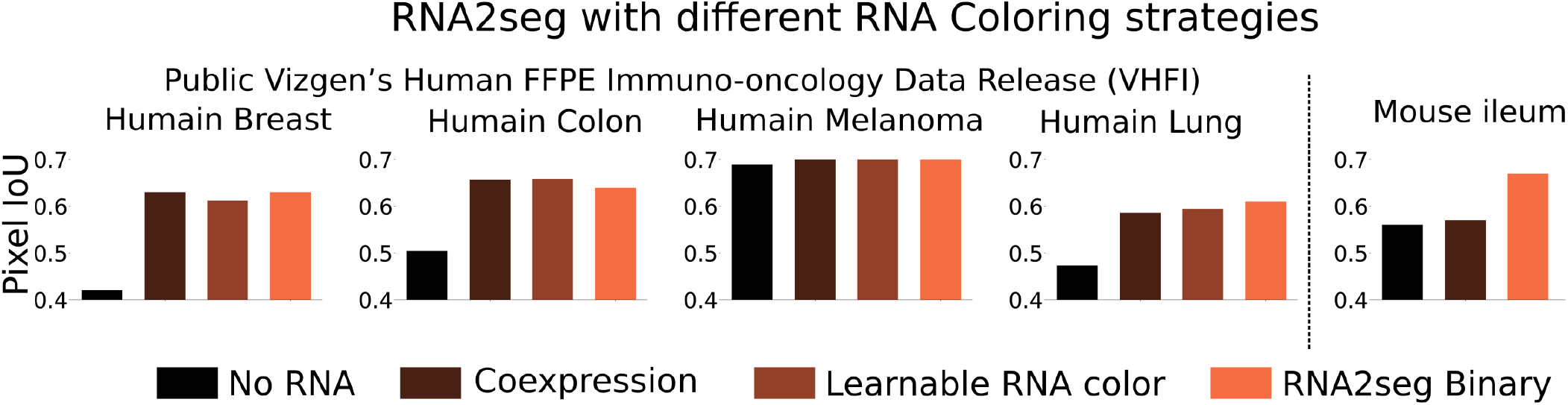
Benchmark of RNA2seg performances using different RNA representations strategies. All models are trained on the VHFI datasets having the same 500 genes spatially resolved. No RNA: RNA2seg trained solely using stainings. Co-expression: RNA2seg trained on RNA images using our co-expression-based coloring method. Learnable RNA Color: RNA2seg where the color of each RNA species is treated as a learnable parameter during training. However, this approach is not transferable to new datasets and is therefore inapplicable to the mouse ileum in a zero-shot setting. Binary: The RNA2seg version retained in this study, trained using only positional information.

**Fig. 5.**
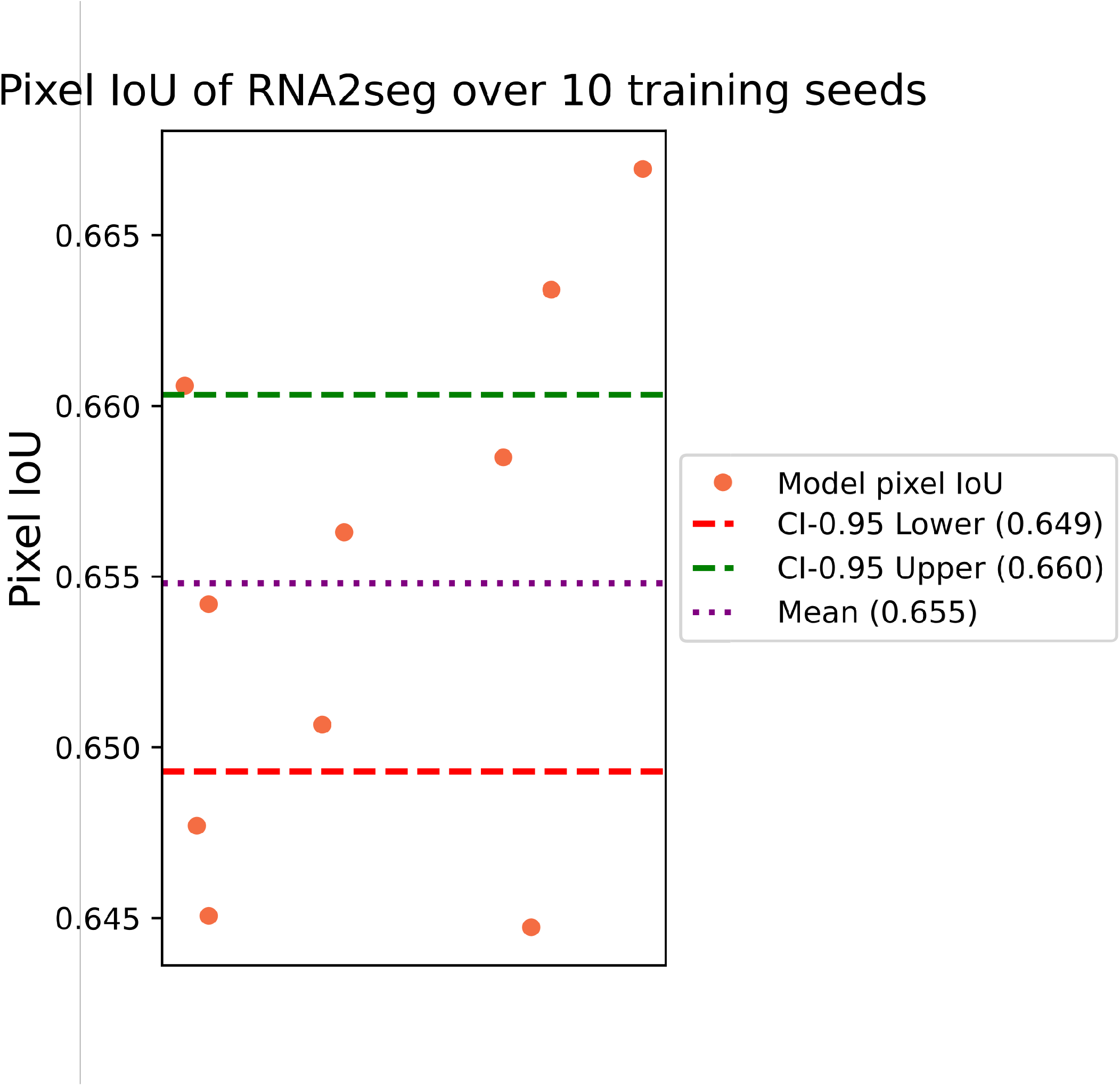
Pixel IoU of RNA2seg-CNN averaged over 10 independent training initializations on all annotated data from the VHFI datasets. The confidence interval (CI) is computed with a t-distribution

